## Supplemental Figure 1-8 + Table 1 for "The TMPRSS2 non-protease domains regulating SARS-CoV-2 Spike in mediated virus entry"

**Supplementary figures**

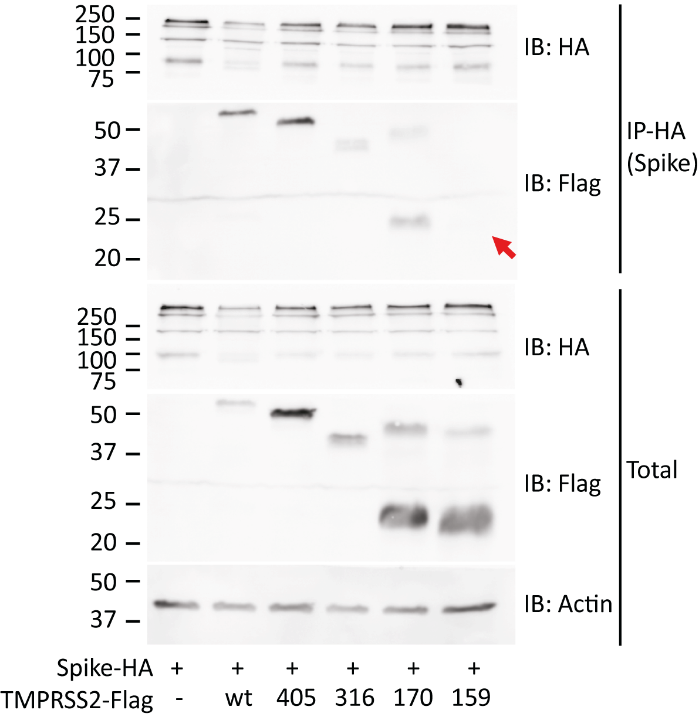

**Figure S1: The reciprocal experiment from WB fig 3.1d** Mapping the Spike-TMPRSS2 interacting region. HEK293T were transfected with the indicated plasmids and IPed with HA-beads and WB were performed as above. Red arrow shows the expected location of the 1-159-TMPRSS2-Flag band within IP-Flag blot.

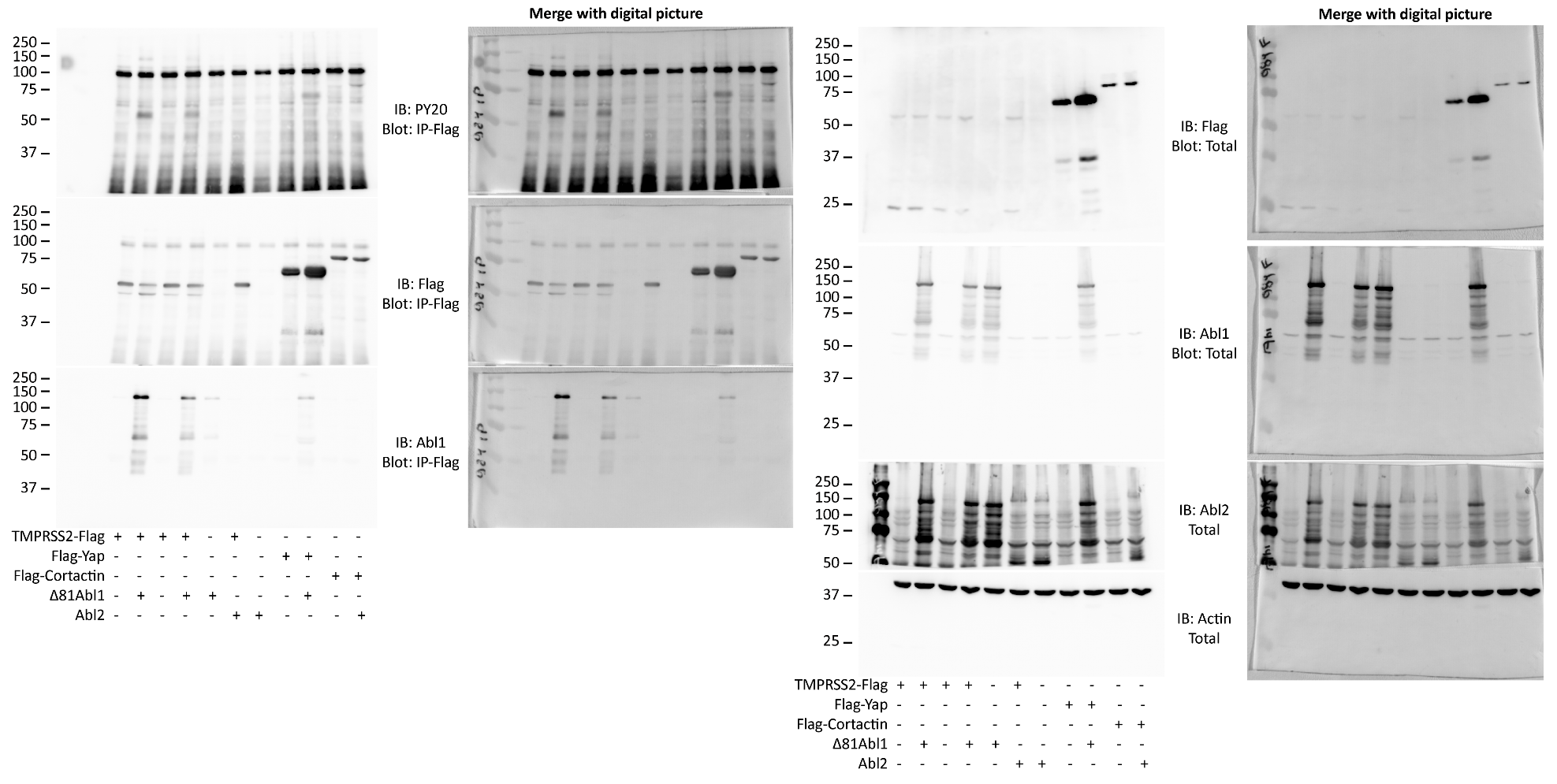

**Figure S2: Original western blot image from Flag-IP figure 1b.** Blots were incubated and re-stained in following order for Flag-IP blot: 1) Anti-mouse-PY20, 2) Anti-mouse-Flag (TMPRSS2, Yap or Cortactin), 3) Anti-mouse-Abl1 and for the total blot: 1) Anti-mouse-Flag (TMPRSS2, Yap or Cortactin), 2) Anti-mouse-Abl1, 3) 2) Anti-mouse-Abl2 and 4) Anti-mouse-Actin

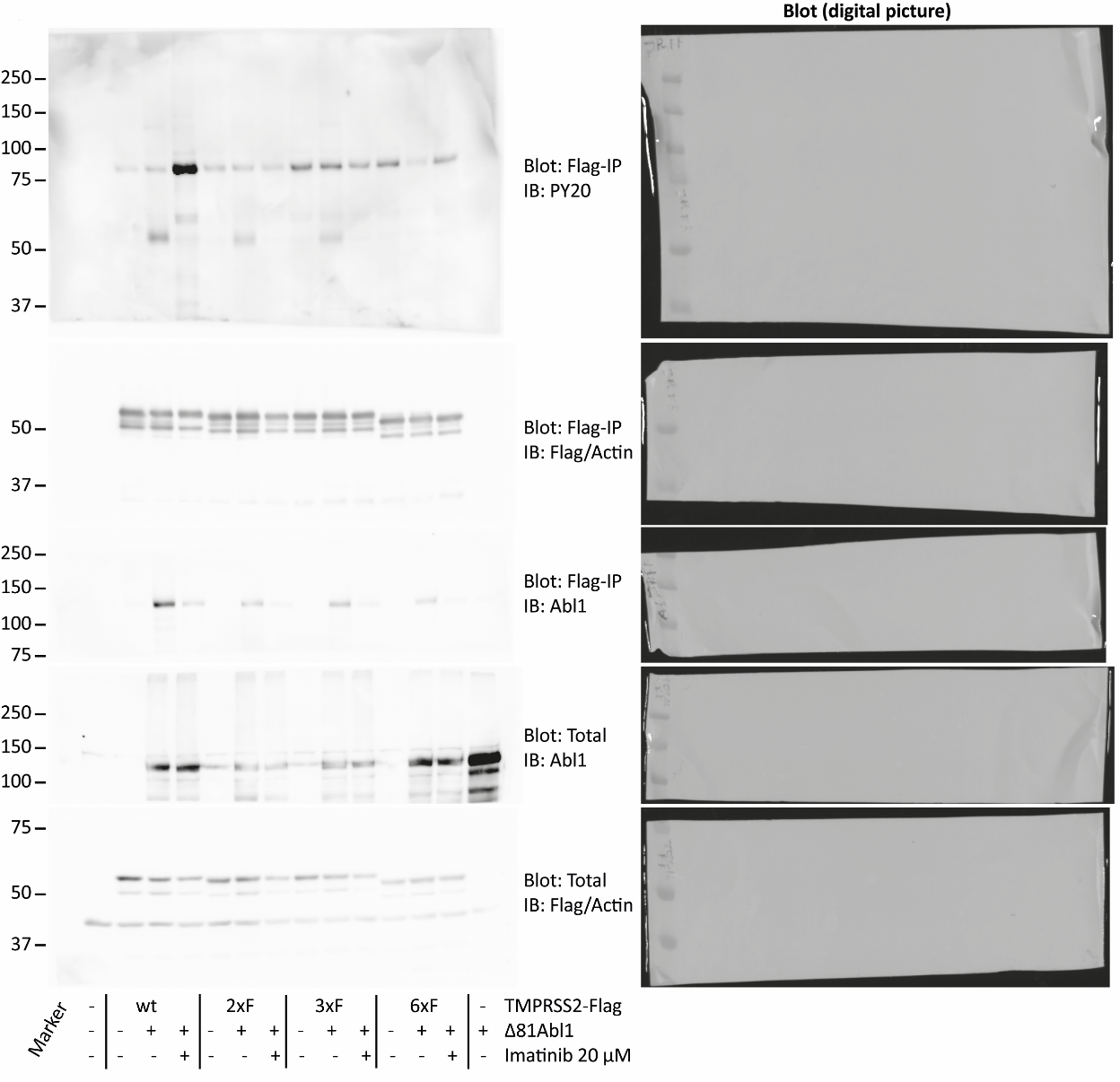

**Figure S3: Original western blot image from Flag-IP figure 1c.** Blots were cut between lane 75 kDa and treated with indicated antibodies. The Flag-IP blot was cut after analysis of PY20 antibody treatment.

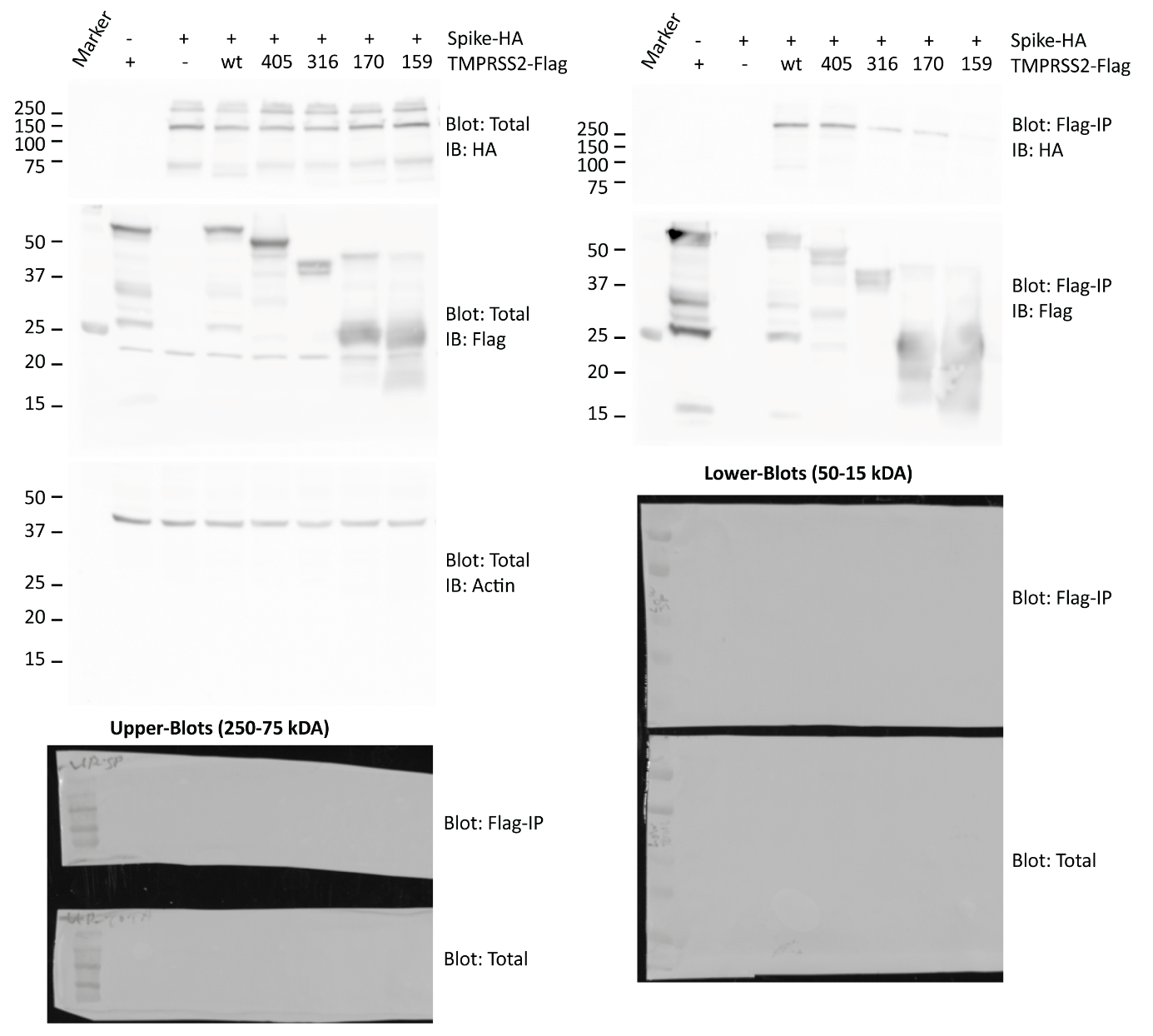

**Figure S4: Original western blot image from Flag-IP figure 2b.** Blots were cut between lane 75 kDa and the upper blots were treated with Anti-mouse-HA (TMPRSS2) antibodies while lower blots was treated with Anti-rabbit-Flag (Spike) and two days later with Anti-mouse-Actin antibodies.

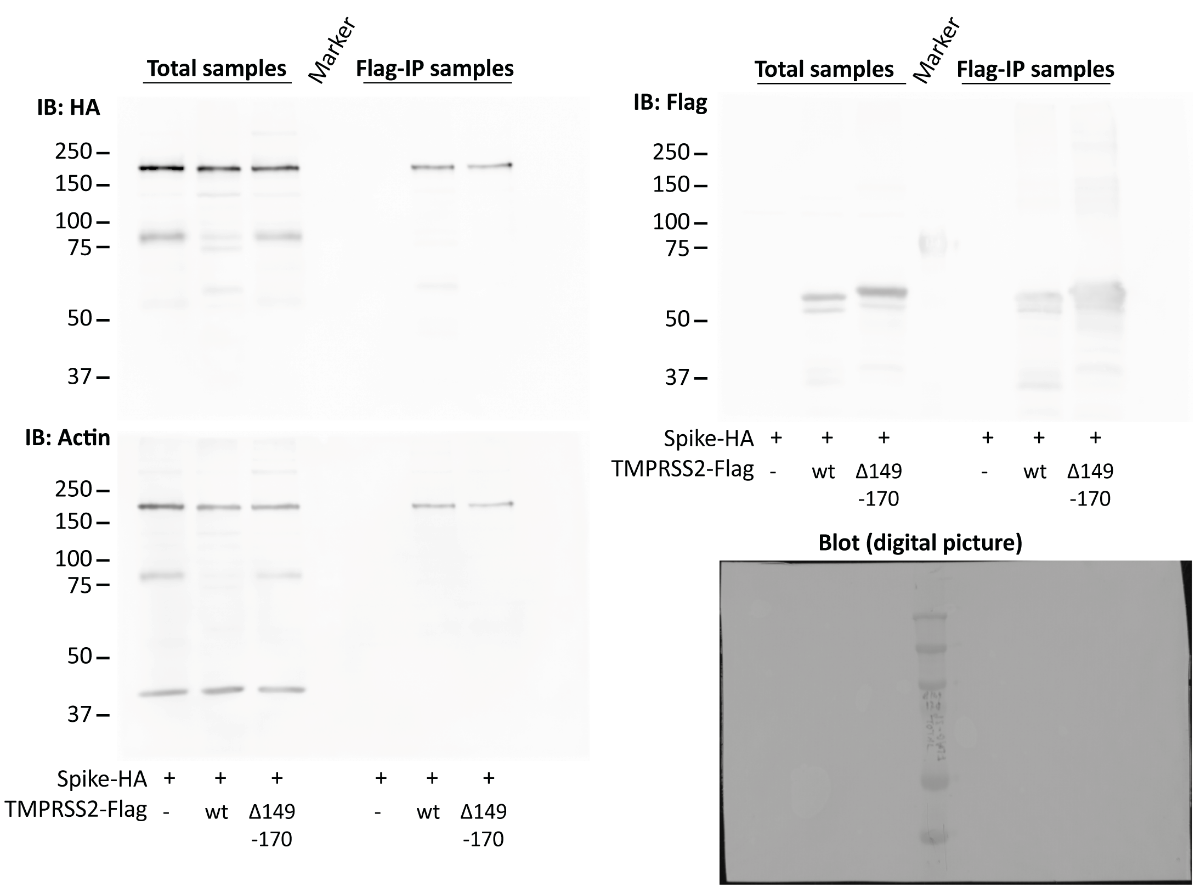

**Figure S5: Original western blot image from Flag-IP figure 2d.** Blot was incubated and re-stained in following order: 1) Anti-mouse-HA (Spike), 2) Anti-rabbit-Flag (TMPRSS2) and 3) Anti-mouse-Actin.

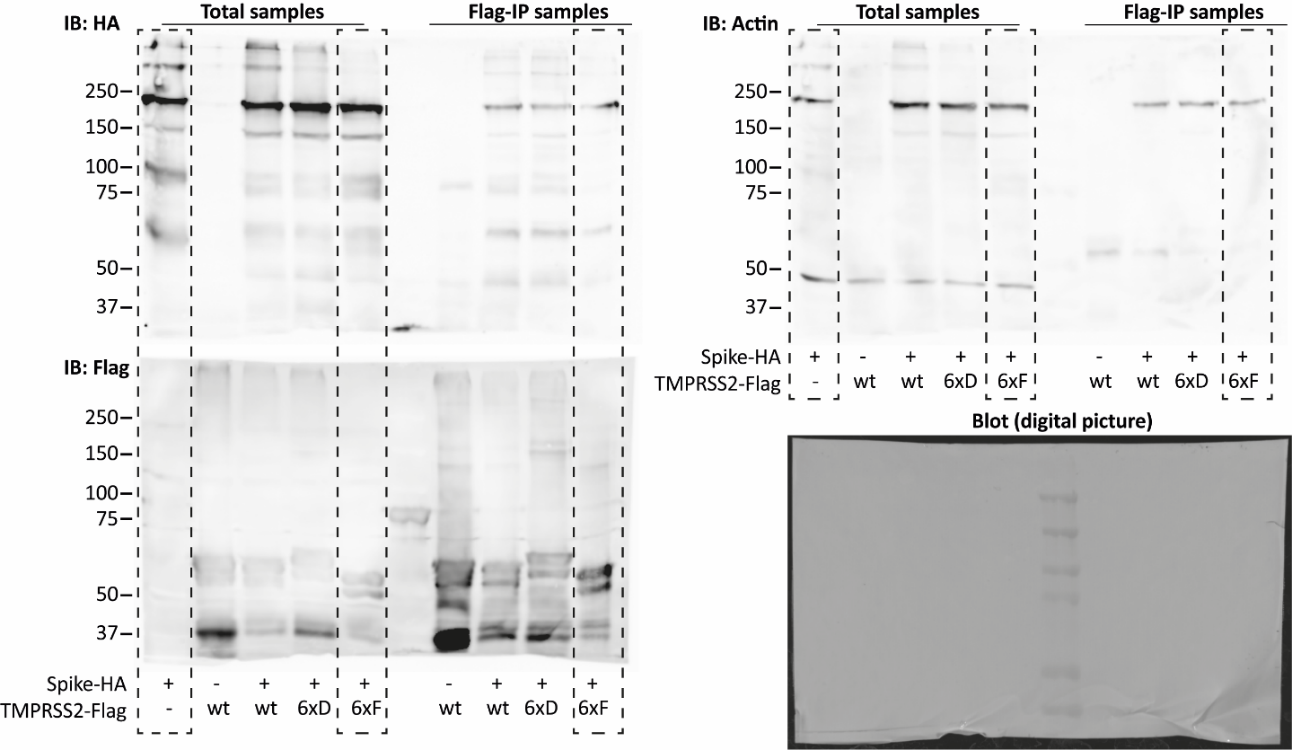

**Figure S6: Original western blot image from Flag-IP figure 4a.** Dashed box shows excluded lanes from figure 4a in main text. Blots were incubated and re-stained in following order: 1) Anti-mouse-HA (Spike), 2) Anti-rabbit-Flag (TMPRSS2) and 3) Anti-mouse-Actin.

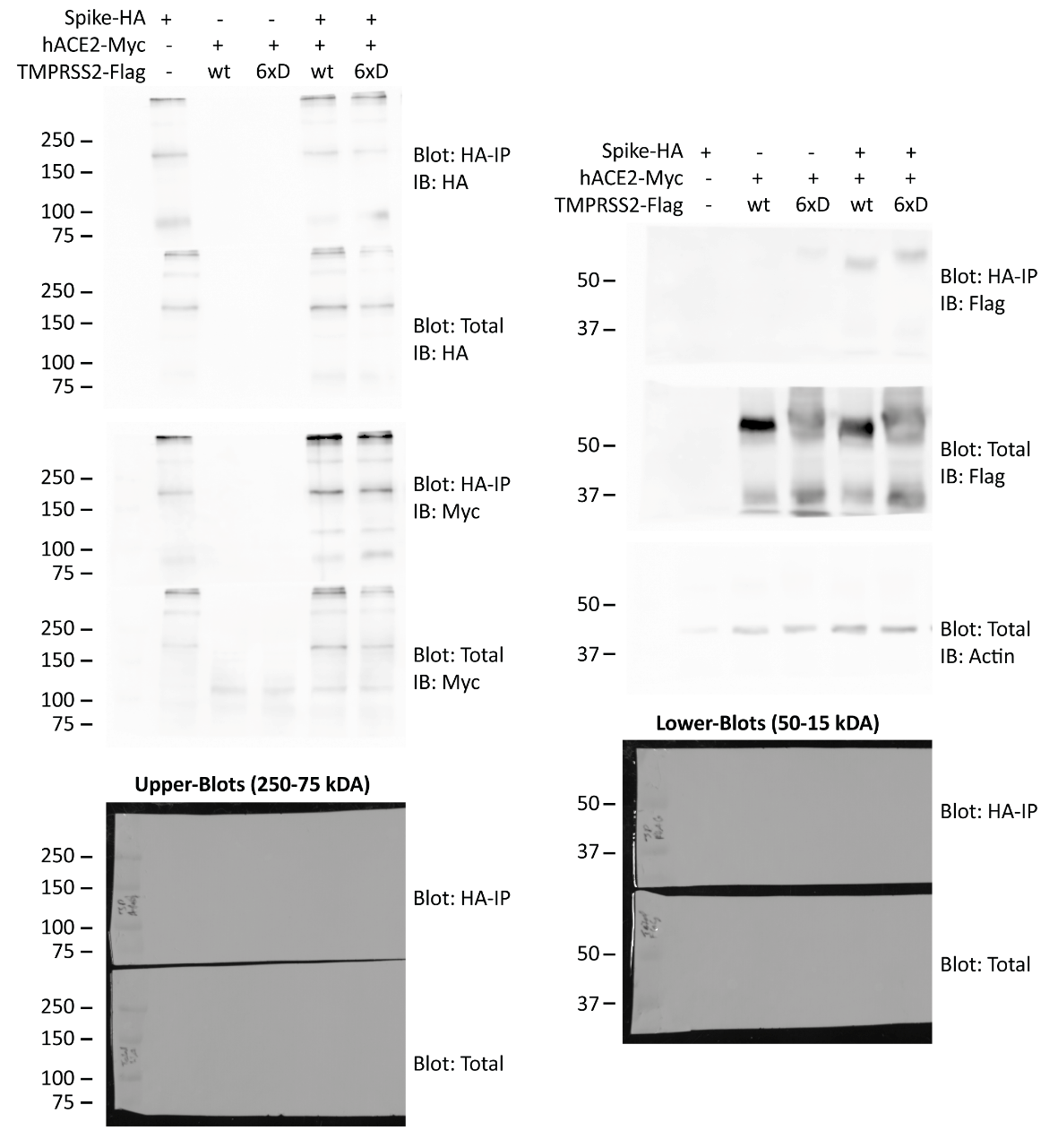

**Figure S7: Original western blot image from Flag-IP figure 4c.** Blots were cut between lane 75 kDa and the upper blots were treated with Anti-mouse-HA (Spike) and anti-Myc (hACE2) antibodies while lower blots was treated with Anti-rabbit-Flag (TMPRSS2) and with Anti-mouse-Actin antibodies. Result was verified with an additional experiment.

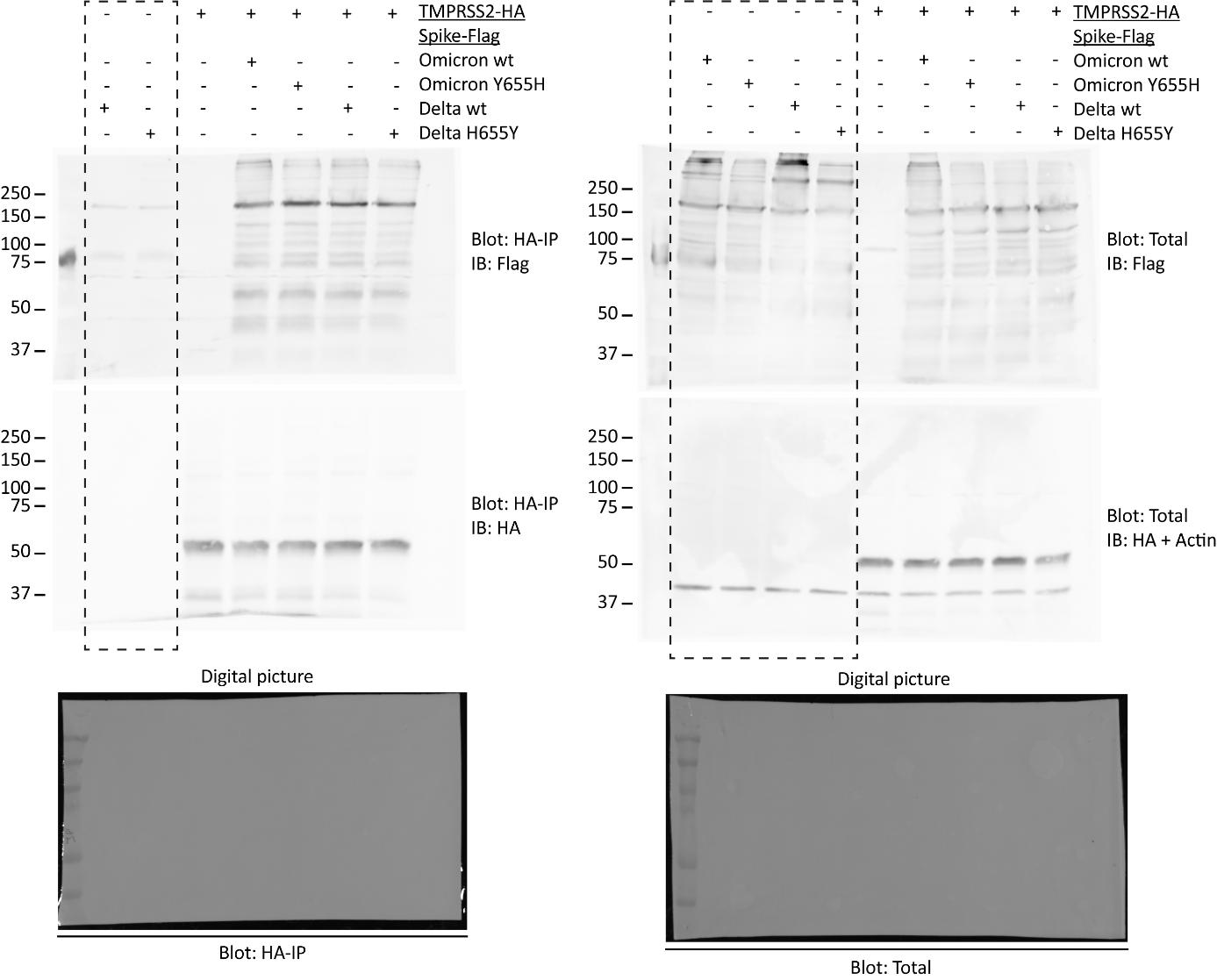

**Figure S8: Original western blot image from Flag-IP figure 5b.** Dashed box shows excluded lanes from figure 5b in main text. Blots were incubated and re-stained in following order: 1) Anti-rabbit-Flag (Spike), 2) Anti-mouse-HA (TMPRSS2) + Anti-mouse-Actin.

**Supplementary tables**

| Name | Sequence |
| --- | --- |
| TMPRSS2_Fw_NheI | AATTGCTAGCGGGCCCACCATGGCTTTGAACTCAGGGTCACC |
| TMPRSS2_Rv_Flag_XbaI | AATTTCTAGATTACTTGTCGTCATCGTCTTTGTAGTCGCCGTCTGCC  CTCATTTGTCGATA |
| TMPRSS2_Rv_HA-tag_XbaI | AATTTCTAGATTAtgcataatccggaacatcatacggataGCCGTCTGCCCT  CATTTGTCGATA |
| TMPRSS2-1347_Rv_Flag_XbaI | CCAATTTCTAGATTACTTGTCGTCATCGTCTTTGTAGTCCTTCGAAGT  GACCAGAGG |
| TMPRSS2-1215_Rv_Flag_XbaI | CCAATTTCTAGATTACTTGTCGTCATCGTCTTTGTAGTCAATGAgAAG  CAccTTGGC |
| TMPRSS2-1083_Rv_Flag_XbaI | CCAATTTCTAGATTACTTGTCGTCATCGTCTTTGTAGTCCACTAGGTC  GTTGAAAGT |
| TMPRSS2-948_Rv_Flag_XbaI | CCAATTTCTAGATTACTTGTCGTCATCGTCTTTGTAGTCTCTCAAAAT  CCCCGCAAA |
| TMPRSS2-513_Rv_Flag_XbaI | CCAATTTCTAGATTACTTGTCGTCATCGTCTTTGTAGTCAGGGTGCC  AGGACTTCCT |
| TMPRSS2-477_Rv_Flag_XbaI | GGGCCCTCTAGATTACTTGTCGTCATCGTCTTTGTAGTCCTGAAGG  ATGAAGTTTGG |
| TMPRSS2-444_Rv_Flag_XbaI | GGGCCCTCTAGATTACTTGTCGTCATCGTCTTTGTAGTCACACCGAT  TCTCGTCCTC |
| TMPRSS2-411_Rv_Flag_XbaI | CCAATTTCTAGATTACTTGTCGTCATCGTCTTTGTAGTCTGACACGC  CATCACACCA |
| TMPRSS2_dAA149-170_Fw | GGACGAGAATCGGTGTGTGTGCCAAGACGACT |
| TMPRSS2_dAA149-170_Rv | AGTCGTCTTGGCACACACACCGATTCTCGTCC |
| TMPRSS2_Rv-YY-FF_P-dead | CACGGGGGACGGGaAGaACTGAGCCGGATG |
| TMPRSS2_Fw-YY-FF_P-dead | CATCCGGCTCAGTtCTtCCCGTCCCCCGTG |
| TMPRSS2_Rv-YY-DD_P-mim | ACGGGGGACGGGTcGTcCTGAGCCGGATGCAC |
| TMPRSS2_Fw-YY-DD_P-mim | TGCATCCGGCTCAGgACgACCCGTCCCCCG |
| TMPRSS2-Y52_Fw-Y-F | CCCGTGCCCCAGTtCGCCCCGAGGGTCCTG |
| TMPRSS2-Y52_Rv-Y-F | GACCCTCGGGGCGaACTGGGGCACGGGGGA |
| 6xf-TMPRSS2-Fw_NheI | CTATAGgctagcGGGCCCACCATGGCTTTGAACTCAGGGTCACCAC  CAGCTATTGGACCTTtCTtTGAAAACCATGGA |
| 6xf-TMPRSS2-Rv | GACCCTCGGGGCGaACTGGGGCACGGGGGACGGGAAGAACTGA  GCCGGATGCACCTCGaAGACAGTGGGGAC |
| 6xd-TMPRSS2-Fw_NheI | CTATAGgctagcGGGCCCACCATGGCTTTGAACTCAGGGTCACCAC  CAGCTATTGGACCTgACgATGAAAACCATGGA |
| 6xd-TMPRSS2-Rv | GACCCTCGGGGCGTcCTGGGGCACGGGGGACGGGTcGTcCTGAGC  CGGATGCACCTCGTcGACAGTGGGGAC |
| Spike_Fw_NheI_pcDNA3.1 | aagctggctagcatgttcgtgttcct |
| Spike-Flag_Rv_XhoI_pcDNA3.1 | ATGCCCctcgagTCActtgtcgtcatcgtctttgtagtctgtatagtGcagtttgacg  cccttc |

**Supplement Table 1: List of Primer used for cloning of described constructs**
